# Single-Cell Proteomics Study of Immune Cell Diversity by Quantitating 465 Proteins

**DOI:** 10.1101/2022.01.07.475463

**Authors:** Liwei Yang, Jesse Liu, Revanth Reddy, Jun Wang

## Abstract

The identification and characterization of T cell subpopulations is critical to reveal cell development throughout life and immune responses to environmental factors. Next-generation sequencing technologies have dramatically advanced the single-cell genomics and transcriptomics for T cell classification. However, gene expression is often not correlated with protein expression, and immunotyping is mostly accepted in the protein format. Current single-cell proteomics technologies are either limited in multiplex capacity or not sensitive enough to detect the critical functional proteins. Herein we present a cyclic multiplex *in situ* tagging (Cyclic MIST) technology to simultaneously measure 465 proteins, a scale of >10 times than similar technologies, in single cells. Such a high multiplexity is achieved by reiterative staining of the single cells coupled with a MIST array for detection. This technology has been thoroughly validated through comparison with flow cytometry and fluorescence immunostaining techniques. Both THP1 and CD4+ T cells are analyzed by the Cyclic MIST technology, and over 300 surface markers have been profiled to classify the subpopulations. This represents the most comprehensive mapping of the diversity of immune cells at the protein level. With additional information from intracellular proteins of the same single cells, our technology can potentially facilitate mechanistic studies of immune responses, particularly cytokine storm that results in sepsis.

## Introduction

Human T cells play a fundamental role in the establishment and maintenance of adaptive immunity^1, 2^. T cells are highly heterogeneous in human adults, since naive T cells have stem cell-like properties and can differentiate into virtually all different types of T cells with specialized functions^3, 4^. For example, they express different receptors with the potential to recognize various antigens from pathogens, tumors, and the environmental stimulations and subsequently generate highly diverse populations such as cytotoxic, memory, effector, or regulatory T cells^5^. In addition, T cell subpopulations are also associated with the life stages and tissue compartment, and different sites exhibit distinct kinetics of changes as individuals age^6, 7^. A well-established phenotypic and functional profiling of T cells would highly contribute to dissect the immunity alterations related to the processes of autoimmune diseases and cancers development and also reveal how immune homeostasis are maintained in response to the increasing number of different pathogens^5, 6, 8^.

With the dramatic progress of single-cell genomics, epigenomics, and transcriptomics, significant advances have been made in methods to annotate T cell subpopulation and the identification of new cell types by single-cell RNA sequencing (scRNA-seq)^9,10, 11, 12, 13, 14^ Recent large-scale efforts, such as the Human Cell Atlas (HCA) project, are attempting to use scRNA-seq to produce cellular maps of entire T cell lineages in human thymus by conducting immunophenotyping at the single-cell level^12^. However, proteins, instead of mRNAs, carry most of the biological functions within a cell and participate in almost all cellular processes including signal transduction, transportation and secretion, di□erentiation and proliferation^15, 16, 17^. For instance, the subsets of memory CD4^+^ and CD8^+^ T cells in peripheral blood mononuclear cells are usually identified based on their profile of secreted cytokines and the expression of cell surface markers like CCR7 and CD62L^18^. Furthermore, protein abundance cannot necessarily be inferred directly from mRNA abundance^19, 20^. The human genome has been estimated to contain approximately 20,000 protein-coding genes, but the number in proteome is considerably higher, as a result of DNA sequence variations, alternative mRNA splicing and posttranslational modifications ^16, 21, 22^. This has been validated by Peterson et al that the CD4 and CD8 proteins of T cells and mRNA copy numbers are highly uncorrelated^24^. Another representative example is that some forkhead box P3 (FOXP3)^+^ regulatory T cells lose FOXP3 gene expression and take on an effector memory T cell phenotype producing interferonγ (IFN-γ)^25, 26^. Therefore, the definition and classification of T cell subpopulation only by the genetic information has been somewhat arbitrary and insufficient, and to fully understand functional properties of cells, those molecular genetic techniques need to be complemented with high performance single-cell protein analyses to define T cell phenotypes^16, 23, 24^.

Current methods for single-cell protein detection are commonly suffered from throughput and multiplex capability^16, 27^, and the range of attainable information normally only span dozens of proteins per cell, which are limited by spectral overlap of fluorophore labels in fluorescence flow cytometry (~17) and the available number of stable metal isotopes in mass cytometry (~40)^28, 29^. Emerging CyCIF (~60) and REAP-seq (~82) techniques increased the detection multiplexity to some extent^24, 30^, but they are still not su cient to cover the whole spectrum of the functional proteome. In this report, we described a single cell cyclic multiplexed *in situ* tagging (Cyclic MIST) technique that are capable of analyzing hundreds to thousands of protein targets with high throughput and high sensitivity. The high content of protein profiling is achieved by multi-round immunostaining with photocleavable oligo barcoded-antibody conjugates and multi-cycle decoding process on complementary oligo modified-microbeads array. Single cells in a microwell chip were reiteratively stained by different conjugates cocktail, with up to 50 conjugates in each staining round. The released oligos from the conjugates upon UV exposure were captured on the MIST array, and the 50 types of captured oligos on each microbead were identified by the decoding process. By repeating the staining round for 10 times, the detection multiplexity by the Cyclic MIST could expand to a level of 465 proteins, a task which is unachievable by current single-cell proteomic techniques. We have thoroughly validated the Cyclic MIST technology and compared its performance with standard flow cytometry and immunofluorescence microscopy using human monocyte THP-1 cells. Most cluster of differentiation (CD) proteins are included in the large panel, and additional 146 intracellular proteins are also profiled simultaneously for every cell. Application of this technology to CD4^+^ T cells leads to discovery of substantially different subpopulations as well as the corresponding intracellular networks pertinent to them. We expect this advanced single-cell proteomic technique could comprehensively profile the diversity of various T cells and provide a cell annotation reference in the identification and characterization of novel or rare T cell types, in addition to providing insights into the underlying mechanisms of T cell development and developing targeted strategies to modulate T cell-mediated immunity in vaccines and immunotherapies.

## Results and Discussion

### Principle of Cyclic MIST Technology

The cyclic MIST platform is a highly multiplexed single-cell proteomics technology which involves multiround antibody staining/elution and multi-cycle decoding procedure (**Figure 1**). Specifically, single cells were loaded into a PDMS microwell chip and simultaneously stained by a cocktail solution of photocleavable oligo barcoded-antibody conjugates. Each antibody was tagged with its unique short oligo by two photocleavable linkers, and one end of the oligo has a biotin moiety. The stained cells were then sealed with a MIST array, which was featured with a highly compact monolayer of oligo-functionalized microbeads. The oligos on the microbeads were complementary to the oligos of the conjugates so that each type of microbeads was able to detect one species of protein from the conjugates cocktail. Upon UV exposure, the released oligos from the single cells were captured and detected on the MIST array by adding streptavidin dye. The signal intensities on the microbeads, which correspond to the amount of released oligos, were proportional to the amount of proteins being detected. Meanwhile, the antibodies on the cells were removed by an elution buffer to recover the antigens, initiating the next staining round by another different panel of conjugates cocktail for cellular targeting.

**Figure 1.**
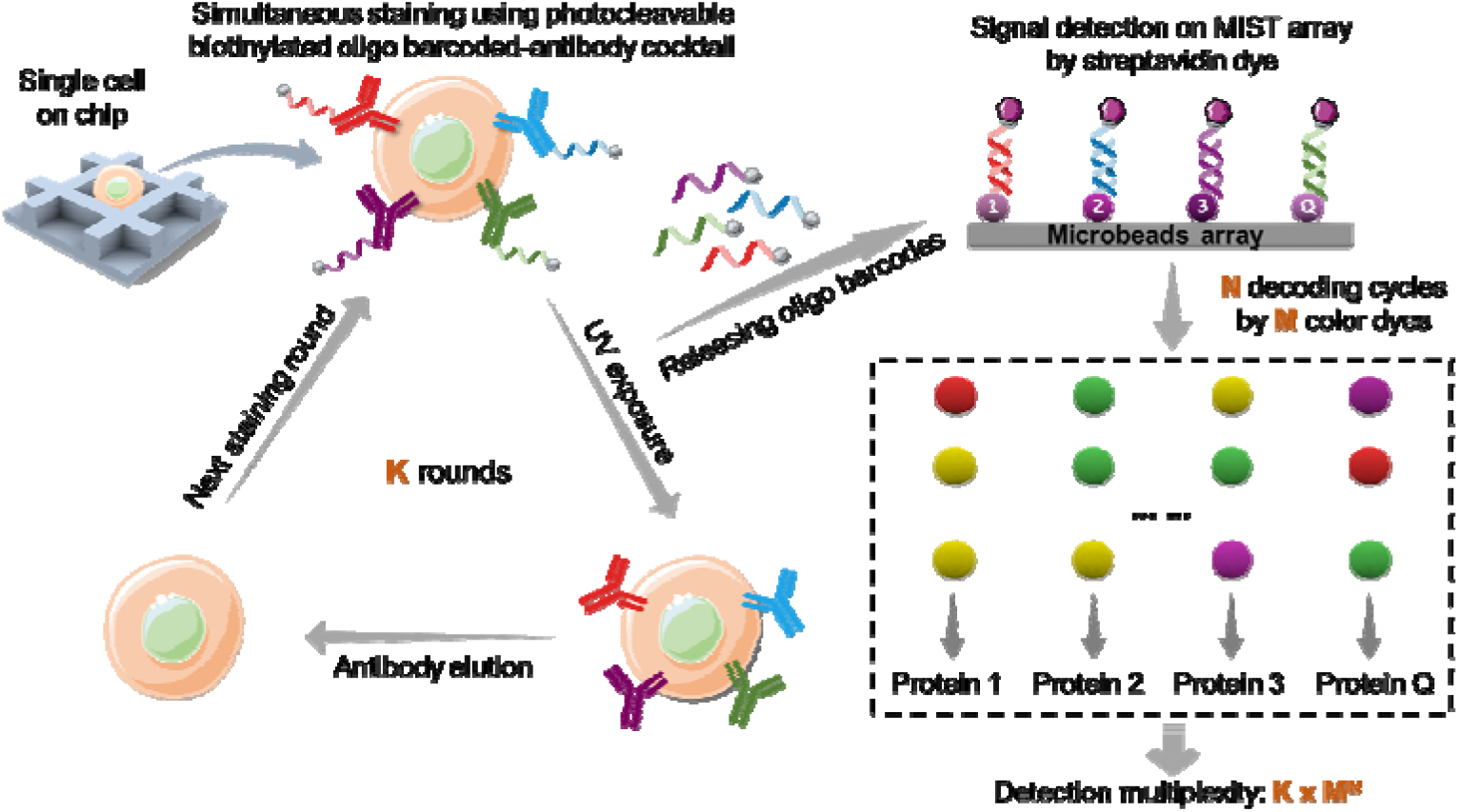
Workflow of cyclic MIST platform for highly multiplexed single immune cell protein analysis. The single immune cells on a PDMS microchip are processed through ***K*** rounds of iterative immunostaining by photocleavable biotinylated oligo barcoded-antibody cocktail. After each round of staining, the released oligo barcodes upon UV exposure are detected on MIST microbeads array. The identities of each microbeads can be assigned by performing ***N*** decoding cycles using ***M*** different colors of complementary oligo-dyes, thus yielding an unprecedented profiling level of protein detection derived as ***K × M^N^***.

To determine which protein/oligo is detected on individual microbeads, a decoding process was conducted through reiterative labeling/quenching cycles of the microbeads. In principle, although all the microbeads are randomly distributed on the whole array, the location of each microbeads is spatially fixed, thus the oligo type on a specific microbead could be identified by successive cycles of dissociation and hybridization process with different fluorophore tagged complementary oligos. After the decoding procedure, each microbead exhibits its particular footprints of color changing, as predesigned in the decoding map, so that the type of oligo or the detected protein by this specific microbead was identified accordingly. Theoretically, performing *N* decoding cycles using *M* different colors of complementary-oligo dyes, in combination with *K* round of cellular targeting, yielding an unprecedented profiling level of protein detection derived as *K × M^N^*. In this report, up to 50 proteins were detected in each staining round to avoid antibodies overcrowding, and 3 decoding cycles by 4 different dyes (Alexa Fluor 488, Cy3, Cy5 and Alexa Fluor 750) were utilized to identify each type of microbeads. After 10 rounds of antibody staining/elution, 465 proteins containing 319 surface marker proteins and 146 intracellular proteins were finally profiled.

### Validation of Cyclic MIST Technology

The cyclic MIST platform was fully validated by using THP-1 human monocyte cell line. We firstly compared cyclic MIST with standard flow cytometry in analysis of membrane protein expression in the THP-1 cells. Five surface antigens (CD107a, CD107b, CD81, HLA-DR and CD147) were measured and analyzed via traditional dot plots by means of the log normalized intensity of the individual protein. For the flow cytometry analysis, the baseline was referred as the isotype-matched controls. For the cyclic MIST technology, the protein intensity from zero-cell microwells (background) plus 3 x standard deviation was considered as the confident threshold. As shown in the **Figure 2a**, both platforms yielded quantitatively similar tendencies for the gated cell populations in dual-proteins analysis, despite that the measured fraction of cells in each quadrant were not exactly matched. For example, the frequencies of CD107b^+^, CD81^+^ and HLA-DR^+^ cells gradually decreased with increasing frequencies of only CD107a^+^ cells by the two techniques. Similarly, the frequencies of only CD107b^+^ cells increased with decreasing frequencies of CD147^+^ and HLA-DR^+^ cells, respectively.

**Figure 2.**
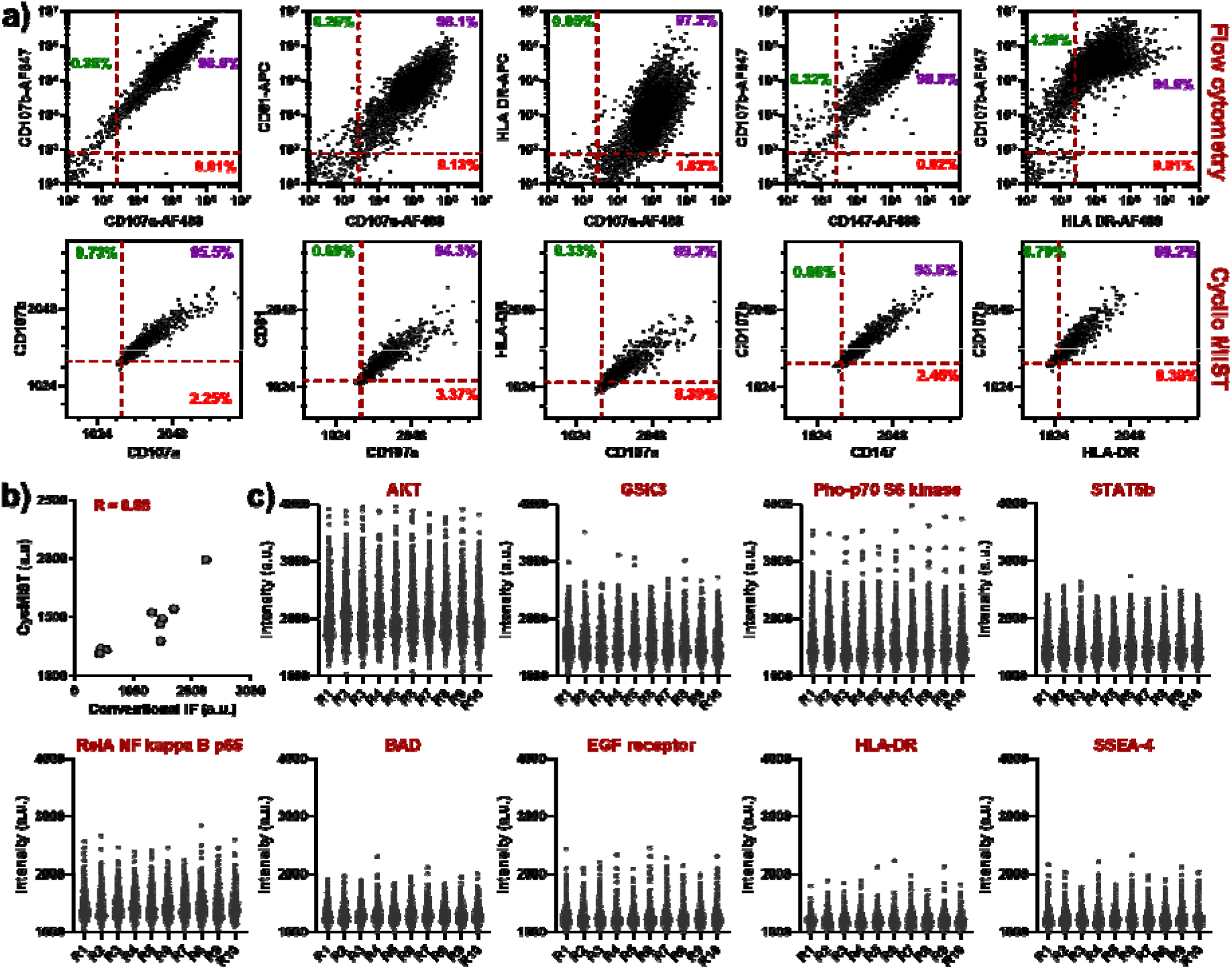
Validation of cyclic MIST platform. a) Representative surface protein-staining results from flow cytometry (n = 20,000 cells) and cyclic MIST (n = 1,512 cells) analysis of THP1 cells. The x and y axis represent the log normalized intensity of the individual protein. For the flow cytometry measurements, the gates were determined by isotype control staining. For the cyclic MIST analysis, the gates separating protein expressed cells and non-protein expressed cells are determined from 0 cell microchip (background) measurements plus 3 x standard deviation. The frequency versus intensity for the individual protein are provided in each quadrant of the scatter plots. b) Correlation of 9 representative protein expression levels between the single cell averages measured by conventional immunofluorescence staining (n =20 cells) and cyclic MIST (n = 400 cells) methods. The correlation coefficient R equals 0.88 for these two methods. c) Representative protein intensities detected by cyclic MIST technology across 10 staining rounds from the same single cells.

We further compared the cyclic MIST measurements to conventional immunofluorescence (IF) staining of bulk assay. Nine representative proteins (AKT, GSK3, Pho-p70 S6 Kinase, STAT5b, RelA/NF kappa B p65, BAD, EGF receptor, HLA-DR and SEEA-4) that covering canonical pathways, cell apoptosis, transcription factor, phosphorylated protein and surface marker proteins were detected by these two techniques. The results were relatively comparable to each other with a correlation coefficient R of 0.88 (**Figure 2b**), despite use of different metrics for assaying expressed protein of cells. The minor discrepancy could be attributed to the different labeling degree of oligos on antibodies and the resultant variation of the antibody-epitope binding. To show the performance of the cyclic MIST for multi-round staining without losing cell integrity, the nine proteins were analyzed across ten staining rounds on the 400 same single cells. The averaged variation of detected signal was calculated as 1.2%, 0.82%, 0.59%, 0.65%, 0.61%, 0.24%, 0.38%, 0.4% and 0.39% for AKT, GSK3, Pho-p70 S6 Kinase, STAT5b, RelA/NF kappa B p65, BAD, EGF receptor, HLA-DR and SEEA-4, respectively, indicating the repeated staining round did not significantly alter the detected protein signals, thus providing the feasibility of analyzing large number of proteins by the cyclic MIST through the expansion of staining rounds.

### Multiplexed Single-Cell Protein Detection by Cyclic MIST Technology

The 465-protein panel has been applied to the THP1 cells and CD4^+^ T cells. The 316 surface markers of the panel include 299 cluster of differentiation (CD) proteins and 17 other membrane proteins. Most known surface markers in hematopoietic cell differentiation are covered by this panel. The additional 146 intracellular proteins involving 55 signaling proteins, 10 transcription factors, and 81 other proteins important to immune response, cell differentiation, and various immune cell functions. 10 rounds of the detection by the cyclic MIST were executed with up to 50 proteins for each. The surface markers are expectedly in lower signal intensities than intracellular proteins in general (**Figure 3a**). The entire population was generally clustered into two groups (**Figure 3b**) on the UMAP figure. That might be because THP1 cells are relatively uniform and have been cultured for almost 10 passages. Through differential expression analysis, we found the cohorts of proteins significantly expressed in both groups (**Figure 3c**). Notably, only the portion of membrane proteins was used in the clustering and differential expression analysis, in order to identify the surface markers for various subpopulations. Gene ontology analysis of the biological process on String indicates that the cluster 0 proteins are more involved in cellcell adhesion and immune response-inhibiting signal transduction. By contrast, the cluster 1 proteins are more related with positive regulation of immune response and cytokine-mediated signaling pathway. Thus, the cluster 0 cells might be more quiescent and tend to adhere to the surface, whereas the cluster 1 cells are receiving certain stimulation that should not be the same simulation by infection since none of them were stimulated in the cell culture. We last examined CD4^+^ T cells with the same panel of 465 proteins by the cyclic MIST technology. After removing the background, the entire population is clustered into 6 groups with distinctive protein expression profiles (**Figure 4a**) where only surface markers are used for clustering. With each group, the proteins show relatively strong correlation with each other (**Figure 4b**). We particularly investigated group 4 and group 3 since their protein profiles are quite different. The dataset allows us to compare intracellular protein expression between these two groups. It was found group 3 cells express more p-BTK, JAK3, cofilin, Ikb-α, T-bet and PTEN proteins than group 4. KEGG pathway analysis indicates that cell differentiation pathways and apoptotic pathways are active in these cells.

**Figure 3.**
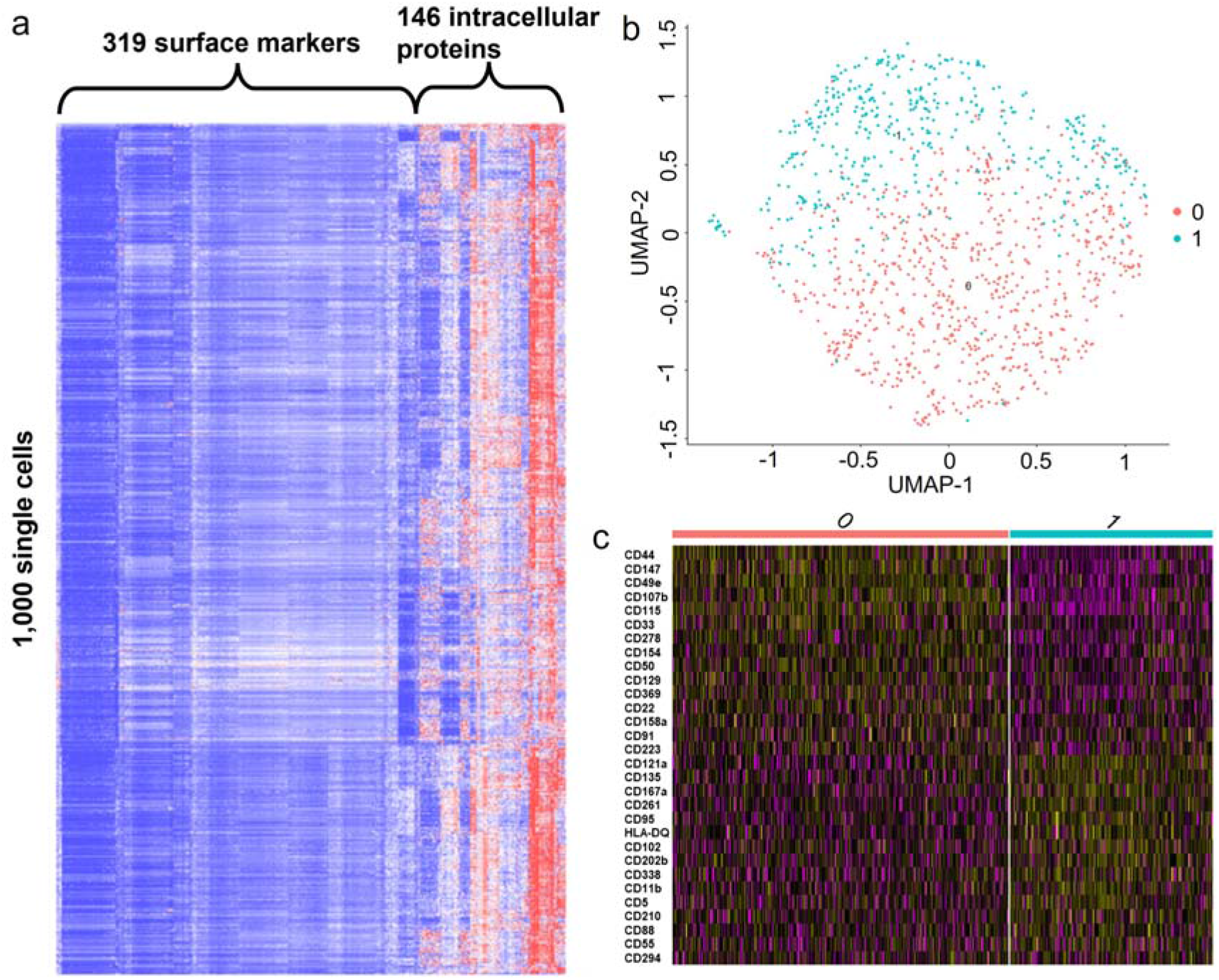
Cyclic MIST assay of single THP1 cells. (a) Heatmap of single-cell expression of 319 surface markers and 146 intracellular proteins for 1000 single cells. (b) UMAP clustering of single cells by surface markers. Two major subpopulations are distinguished. (c) Discovery of differential expression proteins for the two subpopulations shown in heatmap.

**Figure 4.**
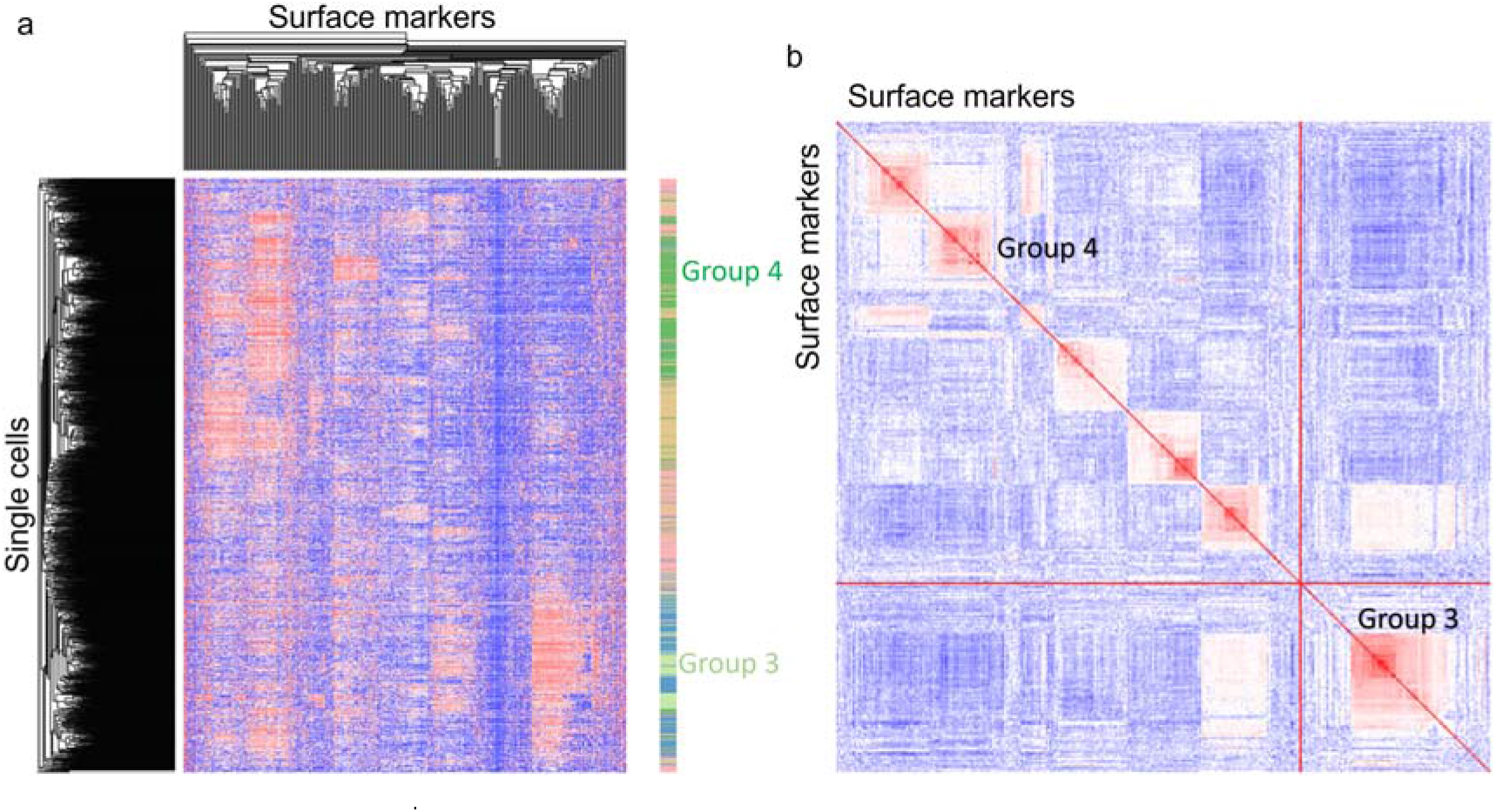
Clustering of CD4^+^ T cells. (a) Heatmap of unsupervised clustering of single cells and surface markers. Only surface markers are used for clustering. 6 groups of subpopulations are identified. (b) Protein-protein correlation heatmap of surface markers. Red indicate relative high values and blue is low values.

## Conclusion

In summary, we report the development of an advanced proteomic technology, Cyclic MIST, to simultaneously measure large number of proteins in single cells. This technique records so far the highest multiplexity (465) for protein analysis in single cells, and the extent of the detection multiplexity could be further improved by simply increasing the immunostaining round and the number of decoding cycles. Comparison of the Cyclic MIST with standard flow cytometry and fluorescence immunostaining demonstrated its high performance in the detection sensitivity, reproducibility, and regenerability of singlecell protein analysis. By applying this technology to CD4^+^ T cells, significant cell heterogeneity and substantially different subpopulations were demonstrated, and further allowing for the analysis of the intracellular networks corresponding to each specific cell subsets. We anticipate the Cyclic MIST could provide new insights in single-cell studies at the protein level to identify phenotypic variation among subpopulations of T cells, and also expect this emerging technique could be coupled with other advanced single-cell sequencing methods to elucidate underlying mechanisms of immune responses and improve the performance of immunotherapy and drug delivery.

## Materials and Methods

### Microchip fabrication

The poly(dimethylsiloxane) (PDMS) microwell chip was fabricated using standard soft-lithographic techniques. A replicate with patterns of photoresists on a silicon wafer was first obtained through photolithography. Then the PDMS prepolymer (Sylgard 184; Dow Corning) and curing agent were mixed in a ratio of 10:1 (w/w) and cast on the patterned replicate. After baking at 80 °C for 2 h, the cured PDMS was separated from the replicate and cut off into pre-designed size and shape for further use. The size of each microwell on the microchip is 50 μm (length) × 50 μm (width) × 40 μm (depth).

### MIST microbeads array preparation

Polystyrene microbeads (2 μm; Life Technologies) were modified with oligonucleotides (15-20 bases) first before patterning. Specifically, 100 μL of amine bearing microbeads solution was firstly reacted with 10 mM bis(sulfosuccinimidyl)suberate crosslinker (BS3; Life Technologies) in phosphate buffered saline (PBS, pH 7.4) solution for 15 mins. After washing by Milli-Q water three times, the microbeads were incubated with 1% poly-L-lysine (PLL; Ted Pella) in pH 8.5 PBS solution for 3 h. The PLL-coated microbeads were rinsed with PBS and further reacted with 300 μM amine-ended oligos (Integrated DNA Technologies) and 2 mM BS3 solutions for 3 h. The microbeads were thoroughly washed with Milli-Q water for further use. Patterning of MIST microbeads array was followed by the previous protocol. Briefly, the 50 types of microbeads carrying different oligos were mixed together in equal portion (16 μL for each) and then mixed with 200 μL of blank microbeads to enhance signal differentiation during imaging. The microbead mixture was deposited onto a cleanroom adhesive tape (VWR) attached on a glass slide and left dry. Subsequently, the array was sonicated for 1 min to remove the excess layers of microbeads on the tape, resulting in the formation of a uniform microbeads monolayer.

### Preparation of photocleavable biotinylated oligo-antibody conjugate

Custom designed biotinylated oligos were purchased from Integrated DNA Technologies and used as received. The crosstalk reactivity of each oligo with other oligos has been validated in our previous report^31^. It is noteworthy that the oligos conjugated to antibodies are complementary to the oligos modified on the microbeads of the MIST array. The conjugation reaction of oligos with antibodies was performed in a dark environment due to the light sensitivity of photocleavable chemicals. Antibodies were concentrated to at least 1 mg/mL using 10 K MWCO centrifuge centrifugal filter (Amicon; Thermo Fisher) and then the solvent changed to a pH 8.0 PBS buffer by 7K MWCO Zeba spin desalting columns (Thermo Fisher). Afterwards, 50 μL of antibodies were reacted with 10 mM photocleavable azido-NHS ester (Click Chemistry Tools) at 1: 25 molar ratio for 2 h. Meanwhile, 30 μL of 200 μM amine-ended biotinylated oligos was incubated with 100 mM photocleavable DBCO-NHS ester (Click Chemistry Tools) at 1: 20 molar ratio for 2 h. The azido-antibodies and DBCO-oligos were both buffer exchanged to pH 7.4 in PBS using the Zeba spin desalting columns and then mixed together for overnight reaction. The resulted conjugates were further purified on a FPLC workstation (AKTA; Bio-Rad) with Superdex^®^ 200 gel filtration column at 0.5 min/mL isocratic flow of PBS buffer. The collected products were concentrated to 0.5 mg/mL using 10 K MWCO centrifuge centrifugal filter and stored at 4°C for further use. The degree of oligo-antibody conjugation efficiency was determined as previously reported^31^.

### Cell culture and processing

The human monocyte cell line THP-1 was purchased from ATCC (TIB-202) and routinely cultured in RPMI 1640 medium (Life Technologies) containing 10% (v/v) fetal bovine serum (FBS), 100 units/mL penicillin, 100 μg/mL streptomycin and 0.05 mM 2-mercaptoethanol (Sigma-Aldrich) at 37 °C in 5% CO_2_ incubator. The THP-1 cells were passaged for 2-3 days with a cell density of 80%-90%. The Human Peripheral Blood CD4^+^ T cells were purchased from Lonza Group Inc. The cryopreserved CD4^+^ T cells were thawed by warming frozen cryovials in a 37°C water bath and transferred into 5 mL of the freshly RPMI 1640 medium. Then the CD4^+^ T cells was centrifugated at 125 × g for 5 mins and washed three times by RPMI 1640, counted using a Countess automated cell counter, and resuspended in RPMI 1640 to a density of 1×10^6^ cells/mL for cell loading in subsequent cyclic MIST assay.

### Flow cytometry analysis

For surface marker staining of THP-1 cells, cells were harvested in 10^6^ cells/mL and split into antibodies and isotype control panels. Then the cells were fixed with 4% paraformaldehyde (PFA) and blocked and permeabilized with a 5% fetal bovine serum (FBS)/0.1% saponin/PBS solution for 30 mins at room temperature (RT). After washing three times with PBS, the cells were incubated with anti-human CD107a, CD107b, CD81, HLA-DR or CD147 conjugated with fluorophore Alexa 488, APC or AF-647 (BioLegend) in the blocking buffer for 1 h at RT. As a negative control, cells were also stained with the control isotype corresponding to each primary antibody. Following a wash step, the cells were resuspended with a 0.5% BSA/PBS solution in a FACS tube and analyzed with a FACSAria IIIu Cell Sorter (BD Biosciences), recording 20,000 events for each sample. The data was analyzed with FlowJo software, and the isotype-matched controls were used as baseline reference for cell segregation.

### Cyclic MIST detection

A cell suspension containing 10,000 cells was applied on a PDMS microwell chip and placed on a shaker with a speed of 40 rpm/min for cell loading. The chip was gently washed with PBS buffer to remove excess cells. After fixation and permeabilization, the cells in the microwells were incubated with a blocking buffer containing 10% goat serum, 1 mg/mL freshly prepared Salmon sperm DNA (Thermo Fisher) and 0.1% Tween 20 in PBS for 1 h. The cells were then stained with a cocktail solution of the photocleavable biotinylated oligos-antibodies conjugates for 1 h. After washing of the cells with PBST containing 0.1% Tween 20 and 5% goat serum, the microchip with cells was mated with a MIST microbead array and clamped tightly. The whole setup was exposed to a 365-nm UV light for 10 mins to release the biotinylated oligos and allow the oligos hybridized on the MIST array. The MIST array was separated from the PDMS chip and washed with 5% goat serum/PBS solution. For protein signal visualization, the MIST array was incubated with 10 μg/mL streptavidin-Alexa Fluor 647 dye (Life Technologies) in 5% goat serum/PBS solution for 15 mins and detected under a fluorescence microscope. Meanwhile, the antibodies on the cells of the microwell chip were washed with a pH 6.6 Gentle Ag/Ab Elution Buffer (Pierce) three time for 5 mins each. Following antibody stripping, cells were rinsed with PBS, washed with the blocking buffer for 5 mins each and blocked for 1 h, initiating the next staining round by adding another conjugate cocktail.

### MIST array decoding process

After signal detection by the microscope, the MIST array was washed by 1 M NaOH solution for 3 mins to dissociate the hybridized oligos, only leaving the single stranded oligos on the microbeads. Subsequently, a cocktail solution of 200 nM complementary oligos tagged with four different fluorophores (Alexa Fluor 488, Cy3, Cy5 and Alexa Fluor 750) in a hybridization buffer (40% formamide and 10% dextran sulfate in saline-sodium citrate buffer) was applied on the array and incubated for 1 h. The array was thoroughly washed by the saline-sodium citrate buffer for three times and then imaged under the fluorescence microscope, obtaining the decoding “Cycle 1” signal. In the second and third decoding cycle, the procedure was followed by the same process as Cycle 1 except that different cocktail solutions of complementary oligos-fluorophores were used, resulting in the decoding “Cycle 2” and “Cycle 3” signal, respectively. These three decoding cycles by four different color dyes can maximally identify 4^3^ (64) types of microbeads/proteins.

### Imaging processing and data requisition

All the images in each array were automatically taken by a Nikon Ti2 inverted fluorescence microscope equipped with a motorized stage, and a lab-developed MATLAB program code was used for image registration, signal quantification, and color determination of each microbeads throughout the three decoding cycles. Basically, the bright field images of microbeads were read first to identify the microbeads’ location, which can be utilized as a reference for image registration. The fluorescence images were used to quantify the intensity of each microbeads on the protein signal and determine the order of fluorescent color change of each microbeads on the decoding signal. After comparing the color order of each microbeads with the protein type in the decoding map, the protein ID of each microbead was determined. Then the microbeads detecting the same protein were compiled together, and their fluorescence intensities were averaged as the intensity of the detected protein on each array. The same procedure was repeated for the whole MIST microbeads array, producing the protein detection signal and their corresponding intensities on each array. After counting the cell number on each microwell of the PDMS chip, a final dataset was generated including the zero or one cell and the detected protein signal/intensity.

### Data analysis and statistics

Raw counts of single-cells and zero-cells from both THP1 and T cells datasets were preprocessed using the Python packages NumPy and pandas^32, 33^. We applied a gating algorithm similar to flow cytometry to preprocess the single-cell data and then removed the batch effect. Single-cell data was first adjusted to remove some background noise in each round by using the equation:

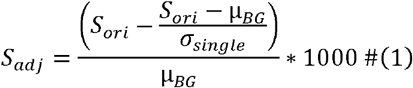

where *S_adj_* is the adjusted signal of the single cells, *S_ori_* is the original signal of the single cells, μ_*BG*_ is the mean of the lowest five values of the microbeads on the MIST array as background level, and *σ_single_* is the standard deviation of the background for each round. The adjusted zero cell data was then used to finish preprocessing the adjusted single-cell data by subtracting each single-cell signal by the mean of the zero-cell protein plus a factor of the standard deviation of the zero-cell protein for that particular signal. Negative values were set to 0 and rows with incomplete data were dropped. The data were further converted in log2 scale before the use of limma’s removeBatchEffect function to remove potential batch effects within the data^34^. The resulting data was then clustered and labeled with their appropriate batch/round group, as a quality control check to ensure that the proteins within the clusters and batches are well dispersed. In both THP1 and CD4+ T cells, the batch corrected datasets were separated into the membrane protein and intracellular protein groups and analyzed using the R package Seurat^35^. The batch corrected datasets were imputed into principal component analysis (PCA) after scaling the data and finding its variable features. Uniform manifold approximation and projection (UMAP) was used for dimension reduction and visualization of the THP1 and T cell clusters after the PCA analysis using the default Louvain algorithm. For the membrane group of the THP1 cell clusters, the UMAP parameters chosen were dims=1:5, n.neighbors=50, spread=2, and min.dist=0.001. For both the intracellular group of the THP1 cell clusters and the membrane group of the T cell clusters, the UMAP parameters chosen were dims=1:15, n.neighbors=50, spread=2, and min.dist=0.001. Both sets of the data were then further analyzed with a heatmap of the top differentially expressed proteins per cluster based on the average log2 fold change within the proteins. Heatmap was either generated by R programs or the Morpheus program (https://software.broadinstitute.org/morpheus). Gene ontology analysis was performed on String with the input of Uniprot IDs for each protein.

## Acknowledgements

This work was supported by the National Institutes of Health R21AG072076 and R01GM128984 allocated to J. W.

## Author contributions

L. Y and J. W designed the study, performed the experiments, provided data analysis and wrote the manuscript. J. L assisted in the data processing and bioinformatic analysis. R. R. optimized the code of MATLAB program. All the authors reviewed the manuscript and approved the submission.

